# Mind the gaps – ignoring errors in long read assemblies critically affects protein prediction

**DOI:** 10.1101/285049

**Authors:** Mick Watson

**Affiliations:** The Roslin Institute and Royal (Dick) School of Veterinary Studies, University of Edinburgh, Easter Bush, EH25 9RG

## Abstract

Long read, single molecule sequencing technologies are now routinely used for whole-genome sequencing and assembly. However, even after multiple rounds of correction, many errors remain which can critically affect protein coding regions, resulting in significantly altered and often truncated protein predictions.

---

Second generation sequencing technologies have revolutionised biological research^1^, largely driven by cheap, high-throughput short-read sequencing technologies^2^. However, it is now clear that these technologies are not suitable for the assembly of large complex genomes, resulting in incomplete and highly fragmented assemblies^3^. Technically and computationally, de novo genome assembly has been considered a “solved problem” for over a decade – one simply needs reads that are longer than the longest repeat region, with sufficient depth and accuracy to detect overlaps between those reads.

Long-read, single-molecule technologies such as those produced by Pacific Biosciences^4^ (PacBio) and Oxford Nanopore^5^ (ONT) have the potential to sequence DNA molecules with lengths in the tens- or hundreds-of thousands of bases, enabling researchers to assemble large and complex repeats. However, both of these technologies have high per-read error rates (in the order of 5-15%), which has resulted in the development of “correction” algorithms. These attempt to use consensus base-calls, raw signal level data and/or shorter more accurate reads to correct long-read assemblies. Examples include Quiver and Arrow^6^ for PacBio, Nanopolish for ONT, and Pilon^7^.

Published genome assemblies from both PacBio^8,9^ and Oxford Nanopore exist^10^. Most recently, Jain *et al* report the genome assembly of a human cell line using ONT’s MinION device, a portable USB sequencer, generating the longest DNA reads ever sequenced. Accuracy after polishing is stated as 99.8% against the current GRCh38 assembly. Accuracy of a recent PacBio-only assembly was reported as 99.7%^9^. By any measure, these are highly accurate assemblies; however, in a genome of over three billion bases, each 0.1% of error represents over 3 million erroneous bases. Moreover, as the predominant errors in both long-read technologies are insertions and deletions, these errors have the potential to critically affect translated regions, which rely on the fidelity of open-reading-frames to predict protein sequences from annotated transcripts.

## Methods

To assess the impact of single molecule assembly errors on protein-coding genes, we downloaded all protein-coding exons from Ensembl, and aligned them to four human reference genomes using BLAT. The genomes chosen were the PacBio assembly from Pendleton *et al*, the ONT assembly from Jain *et al*, the CHM1 genome after two rounds of polishing from Koren *et al*^8^ and the GRCh38 primary assembly from Ensembl. The latter serves as a positive control. Short exons (<300bp) were removed, and alignments only considered where > 90% of the exon was contained within a single alignment.

## Results

The results can be seen in table 1.

**Table 1.** Alignment statistics from a comparison of all protein coding exons (>300bp) against three single-molecule sequencing assemblies and the GRCh38 reference genome

|  |  |  |  |  |
| --- | --- | --- | --- | --- |
| **Total exons** | 73792 |  |  |  |
| **Total genes** | 20932 |  |  |  |
|  | **Exons with full length alignment** | **Total # mismatches** | **Total gap openings** | **# protein-coding genes with gaps** |
| GRCh38 | 73689 | 5082 | 291 | 161 |
| Nanopore (Jain *et al*) | 64481 | 53275 | 18098 | 7022 |
| PacBio1 (Pendleton *et al*) | 69972 | 56601 | 125921 | 16630 |
| PacBio2 (Koren *et al*) | 71563 | 53711 | 11327 | 5272 |

The single molecule assemblies contain a high number of mismatches and gaps. Whilst the three assemblies have a similar number of mismatches, the PacBio assembly from Pendleton *et al* has a far higher number of gaps (indels). The indels in the nanopore assembly are predicted to affect 7022 (33%) of the protein-coding genes considered; the PacBio assembly from Pendleton *et al* 16630 (80%) genes; and the PacBio assembly from Koren *et al* 5272 (25%) of genes. Whilst some of these exons contain UTR sequence, the majority of the mismatches and gaps will fall within protein coding exons and will undoubtedly have a critical and often devastating effect on the resulting protein prediction.

Mismatches and gaps against GRCh38 should be considered an estimate of the error rate inherent in our (simple) alignment protocol. Some of the mismatches and gaps in the ONT and PacBio assemblies will represent genuine differences between the GRCh38 assembly and the actual genome sequenced; however, these are likely to explain only a tiny proportion of the differences observed.

## Discussion

There is a substantial improvement between the PacBio assembly produced by Koren *et al* compared to that produced by Pendleton *et al*. As well as benefitting from improved data quality from more recent sequencing chemistry, the Koren *et al* assembly underwent two rounds of Quiver polishing, as well as correction of the reads during assembly by Canu. Together, these steps may explain the observed improvement. The ONT assembly benefitted from Pilon correction with short Illumina reads as well as raw signal-level polishing. However, many indels remain because of the problems inherent with mapping short Illumina reads to repetitive sequences (which includes gene families). If reads do not map, or map to multiple locations (a known issue in RNA-Seq^11^), then no correction can be performed.

These results should not be considered a criticism of either PacBio or ONT, both of which are highly accurate technologies; nor should they be considered a criticism of Pendleton *et al*, Jain *et al* or Koren *et al*, all of which are ground-breaking pieces of research. Rather, the results indicate that even after multiple rounds of polishing, critical errors remain in single molecule assemblies, that can affect protein predictions, and which only many months of manual effort can fix. This conclusion has ramifications across the biological and medical sciences, for those researchers seeking to sequence genomes (and seek funding to sequence genomes) using single molecule technologies.

A pipeline to reproduce the above analysis can be found at: https://github.com/WatsonLab/sm_assemblies

## References

1. Goodwin, S., Mcpherson, J. D. & Mccombie, W. R. Coming of age: ten years of next-generation sequencing technologies. Nat. Publ. Gr. 17, 333–351 (2016).

2. Watson, M. Illuminating the future of DNA sequencing. Genome Biol. 15, 108 (2014).

3. Bradnam, K. R. et al. Assemblathon 2: evaluating de novo methods of genome assembly in three vertebrate species. Gigascience 2, 10 (2013).

4. Eid, J. et al. Real-time DNA sequencing from single polymerase molecules. Science 323, 133–8 (2009).

5. Loman, N. J. N. J. & Watson, M. Successful test launch for nanopore sequencing. Nat. Methods 12, 303–4 (2015).

6. Chin, C.-S. et al. Nonhybrid, finished microbial genome assemblies from long-read SMRT sequencing data. Nat. Methods 10, 563–569 (2013).

7. Walker, B. J. et al. Pilon: An Integrated Tool for Comprehensive Microbial Variant Detection and Genome Assembly Improvement. PLoS One 9, e112963 (2014).

8. Koren, S. et al. Canu: scalable and accurate long-read assembly via adaptive k-mer weighting and repeat separation. Genome Res. 27, 722–736 (2017).

9. Pendleton, M. et al. Assembly and diploid architecture of an individual human genome via single-molecule technologies. Nat. Methods 12, 780–786 (2015).

10. Jain, M. et al. Nanopore sequencing and assembly of a human genome with ultra-long reads. Nat. Biotechnol. (2018). doi:10.1038/nbt.4060

11. Robert, C. & Watson, M. Errors in RNA-Seq quantification affect genes of relevance to human disease. Genome Biol. 16, 177 (2015).

